# tauX: A Gene Expression Ratio Strategy to Improve Machine Learning Applications in Precision Medicine

**DOI:** 10.1101/2024.07.01.601595

**Authors:** Jacob Pfeil, Liqian Ma, Hin Ching Lo, Tolga Turan, R. Tyler McLaughlin, Xu Shi, Severiano Villarruel, Stephen Wilson, Xi Zhao, Josue Samayoa, Kyle Halliwill

**Affiliations:** GRC Computational Oncology, AbbVie Bay Area, 1000 Gateway Boulevard, South San Francisco, CA 94080; GRC Methods Development, AbbVie Bay Area, 1000 Gateway Boulevard, South San Francisco, CA 94080

**Author notes:** Corresponding author: Jacob Pfeil.

## Abstract

Machine learning algorithms identify patterns that would otherwise be difficult to observe in high-dimensional molecular and clinical data. For this reason, machine learning has the potential to have a profound impact on clinical decision making and drug target discovery. Nevertheless, there remain considerable technical challenges in adapting these tools for clinical use. These challenges include clinical feature engineering, model selection, and defining optimal strategies for model training. For cancer care, RNA sequencing of patient tumor biopsies has already proven to be a powerful molecular assay to characterize tumor-intrinsic and -extrinsic phenotypes influencing therapeutic response. To improve the predictive performance of RNA-sequencing data, we developed the tauX machine learning framework to refine gene expression features and improve the performance of machine learning algorithms. The tauX framework uses aggregated ratios of positively and negatively associated predictive genes to simplify the prediction task. We showed a significant improvement in predictive performance using a large database of synthetic gene expression profiles. We also show how the tauX framework can be used to elucidate the mechanisms of response and resistance to checkpoint blockade therapy using data from the Stand Up to Cancer (SU2C) Lung Response Cohort and The Cancer Genome Atlas (TCGA). The tauX strategy achieved superior predictive performance compared to models built upon established feature engineering strategies or widely used cancer gene expression signatures. The tauX framework is available as a freely deployable docker container (https://hub.docker.com/r/pfeiljx/taux).

## Introduction

While recent advancements in nucleotide sequencing technology and analytical techniques have aided in the development of molecularly informed therapeutic strategies, evidence is accumulating that patient-specific effects strongly influence treatment outcomes (Bretthauer et al., 2023; Zhang et al., 2019). Machine learning/artificial intelligence (ML/AI) approaches have been proposed as a solution to this problem, but despite an era of remarkable development in machine learning and artificial intelligence, there are only a few examples of successful applications of ML to precision medicine and clinical drug development tasks (Shah et al., 2019; Vasey et al., 2021). The reasons for the limited applications include a combination of conceptual, technical, and regulatory hurdles. Examples of challenges outside of the regulatory framework surrounding the clinical use of AI technology include the complexity of biological data and the limited number of clinically relevant training datasets. These challenges are unlikely to be solved by increasingly sophisticated ML algorithms, as more complex models perform poorly on high-dimensional data with a small number of samples (Altman & Krzywinski, 2018).

While genetic data are commonly used for subtyping cancer patients and matching patients to targeted therapies, only a small proportion of patients carry actionable genetic markers that allow them to benefit from this profiling (Rodon et al., 2019). In contrast to the specificity of targeted panels of actionable molecular alterations, whole-transcriptome gene expression data may inform treatment decisions for a broader swath of patients regardless of the presence of specifically targetable alterations. The development of superior predictive signatures derived from whole-transcriptome gene expression data paired with flexible modeling solutions may facilitate the adoption of ML and transcriptome data for clinical decision-making (Morozova et al., 2009; Turan et al., 2021).

Traditional RNA-seq analysis includes differential expression analysis, where RNA-seq counts are modeled as negative binomial distributions and significant differences are determined using a hypothesis test (Anders & Huber, 2010). Differentially expressed genes may be used alone or for gene-set enrichment analysis (GSEA). GSEA is a well-established technique for understanding the coordinated expression of complex biological systems (Barbie et al., 2009; Hänzelmann et al., 2013; Subramanian et al., 2005). Genes known to contribute to specific biological processes have been assembled into gene set databases, including MSigDB, Reactome, KEGG, and the Gene Ontology Knowledgebase (Ashburner et al., 2000; Gillespie et al., 2022; Kanehisa & Goto, 2000; Liberzon et al., 2011; Subramanian et al., 2005). For gene expression analysis of cancer samples, GSEA can measure the relative level of immune infiltration, stromal composition, and tumor intrinsic pathway activation. It has also been shown that the ratio of specific gene sets or other molecular and cellular features can further improve predictions of clinical outcomes, including survival and response to therapies (Sinicrope et al., 2020; Turan et al., 2021). However, a generalizable framework for the optimization of predictive gene expression ratios has not been thoroughly explored.

Here, we describe a computationally intensive gene expression transformation that internally normalizes gene expression profiles using pairwise gene ratios. Using synthetic and clinical lung cancer data, we showed that superior predictive accuracy can be achieved using gene expression ratios. For example, gene ratio features achieved the highest validation score in the recent anti-PD1 response prediction challenge (Mason et al., 2022; Vincent et al., 2021). Further characterization of engineered gene-set ratios revealed new enriched biological pathways involved in the response to checkpoint blockade therapy that could lead to improved patient subtyping and response prediction. This framework makes no assumptions about the prediction task and can be applied to any precision medicine task involving binary outcomes and gene expression data.

## Methods

### Generation of Synthetic Gene Expression Data

Synthetic gene expression data were generated using the TCGA lung adenocarcinoma (LUAD) study (N=601) (Campbell et al., 2016; Collisson et al., 2014). We varied several important parameters to explore the effect of important gene expression and sample subtype parameters. These parameters included the effect size (Cohen’s *d*: 3.0, 2.0, 1.0, 0.75, 0.5, 0.25), number of DEGs (100, 50, 30, 20, 10), percentage of synthetic responders (0.25, 0.2, 0.15, 0.10, 0.05), and strength of the correlation between DEGs (Spearman R: 0.9, 0.75, 0.5, 0.25). The parameter sweep yielded 600 unique experiments and generated 240,000 unique synthetic gene expression profiles for training and testing the tauX framework.

The recount3 recomputed counts for the TCGA LUAD study were used to model the lung adenocarcinoma gene expression distribution (Wilks et al., 2021). Gene expression counts were scaled to transcripts per million (TPM) and normalized using a log2(TPM + 1) transformation (Zhao et al., 2021). The means and covariances for the background distribution were calculated to model the correlation structure of the TCGA-LUAD cohort. Background gene expression profiles were then sampled from the resulting multivariate normal distribution. Responder gene expression profiles were sampled from a modified background distribution where the means were shifted to accommodate the statistical properties for each experiment. Synthetic LUAD gene expression profiles were then used for downstream precision medicine subtyping experiments.

### Traditional Gene Expression Feature Engineering Approaches

The synthetic gene expression profiles were visualized as a hierarchically clustered heatmap using the Ward algorithm (Bar-Joseph et al., 2001; Waskom, 2021). The tauX approach was compared to two routine preprocessing steps for developing gene expression machine learning algorithms. The first is to rank the log-normalized genes by their variance to obtain the most highly variable genes (HVGs) (Lun et al., 2016). The second strategy was to correlate log-normalized gene expression values with the binary response outcome variable using a Z score to obtain the most differentially expressed genes (DEGs) (Cheadle et al., 2003). The HVGs and the DEGs were used as input for training machine learning tools.

### tauX Response Feature Engineering Approach

The tauX strategy uses gene feature counts normalized to transcripts per million (TPM) values. To ensure numerical stability, the TPM values are rescaled using the min–max scaler to a range between 1 and 100 (Pedregosa et al., 2011). Outlier gene expression was also rescaled to 1.5 times the interquartile range value (Tukey & others, 1977). The C++ programming language was used to make the pairwise ratio calculations more efficient. Gene ratios were then compared between responder and nonresponder samples using Student’s t-statistic to identify differentially expressed gene ratios (DEGRs).

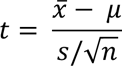

The resulting DEGRs were subsequently clustered to identify modules of gene ratio expression associated with response. Since the input DEGRs were positively correlated with response, the numerator genes were associated with response, and the denominator genes were associated with nonresponse. Specific biological functions were investigated using gene set overlap analysis (Badia-i-Mompel et al., 2022). The mathematical structure of the ratios allowed us to detect enrichment of the ratio associated with expression using the SingScore bidirectional gene set enrichment approach (Foroutan et al., 2018). Specifically, the numerator genes were used as the up signatures, and the denominator genes were used as the down signatures.

### Automated Machine Learning Model Generation

The optuna Bayesian hyperparameter optimization framework was used to train each of the machine learning models (Akiba et al., 2019). Using the optuna framework ensured that the best performing model was used when evaluating each feature engineering approach. The input features from the traditional and tauX feature engineering strategies were used to train commonly used machine learning algorithms, specifically elastic net regression, linear support vector machine (SVM), and radial basis function (RBF)-kernel SVM (Akiba et al., 2019; Pedregosa et al., 2011). In every case, the machine learning model was evaluated using a hold-out set of samples.

### Generation of Stand-Up to Cancer (SU2C) Lung ICI Response Training and Testing Data

Gene expression counts and clinical data for 152 SU2C samples were used to generate training and validation data (Ravi et al., 2023). The SU2C Lung cohort included lung adenocarcinoma (LUAD) and lung squamous cell carcinoma (LUSC) patient samples. The expression data were then compared to those of patients who received anti-PD(L)1 therapy and experienced progressive disease (PD), partial response (PR), or complete response (CR). Outlier gene expression profiles were identified using isolation forests (Liu et al., 2008), local outlier factor analysis (Breunig et al., 2000), and one-class SVM (K.-L. Li et al., 2003). Consensus clustering was used to identify expression subtypes (Lai et al., 2021; Monti et al., 2003; Ott, 2022). The number of clusters was selected by optimizing the Bayesian information criterion, Davies–Bouldin index, silhouette score, and *Calinski–Harabasz* index.

The training and testing cohorts were generated by random sampling (80:20 split), and the anti-PD1 therapy response label and consensus cluster assignments were used to stratify the patients (n=72). The tauX gene expression signatures were learned by applying the tauX approach to the SU2C training data. The resulting ratios were clustered to identify modules of correlated expression and to generate bidirectional gene sets. The SingScore enrichment algorithm was used to stably score single samples without the need for a background cohort (Foroutan et al., 2018). The resulting gene-set enrichment scores were used to train the elastic net, linear SVM, and nonlinear RBF-SVM models.

### Survival analysis

K‒M plots and log-rank statistics were generated using the R survminer package (Kassambara et al., 2021). Cox proportional hazards models were fit using the R survival package (Therneau & Grambsch, 2000). The plots were generated using ggplot (Wickham, 2016) and matplotlib (Hunter, 2007).

## Results

### Overview of the tauX Gene Expression Modeling Framework

The tauX approach was designed to identify paired gene expression effects that correlate with the response to immune checkpoint blockade (Figure 1). However, this approach makes no assumptions about the prediction task and therefore can be applied to many different machine learning tasks. The goal of the tauX modeling framework is to minimize the background noise and maximize the signal associated with clinical response. The combinatorial complexity of this problem was computationally challenging (i.e., C_2_(10,000) ≈ 5 × 10^7^ comparisons). To scale to an entire transcriptome, the computationally efficient C++ programming language was used to accelerate the identification of gene expression ratio features (Supplementary Figure 1).

**Figure 1.**
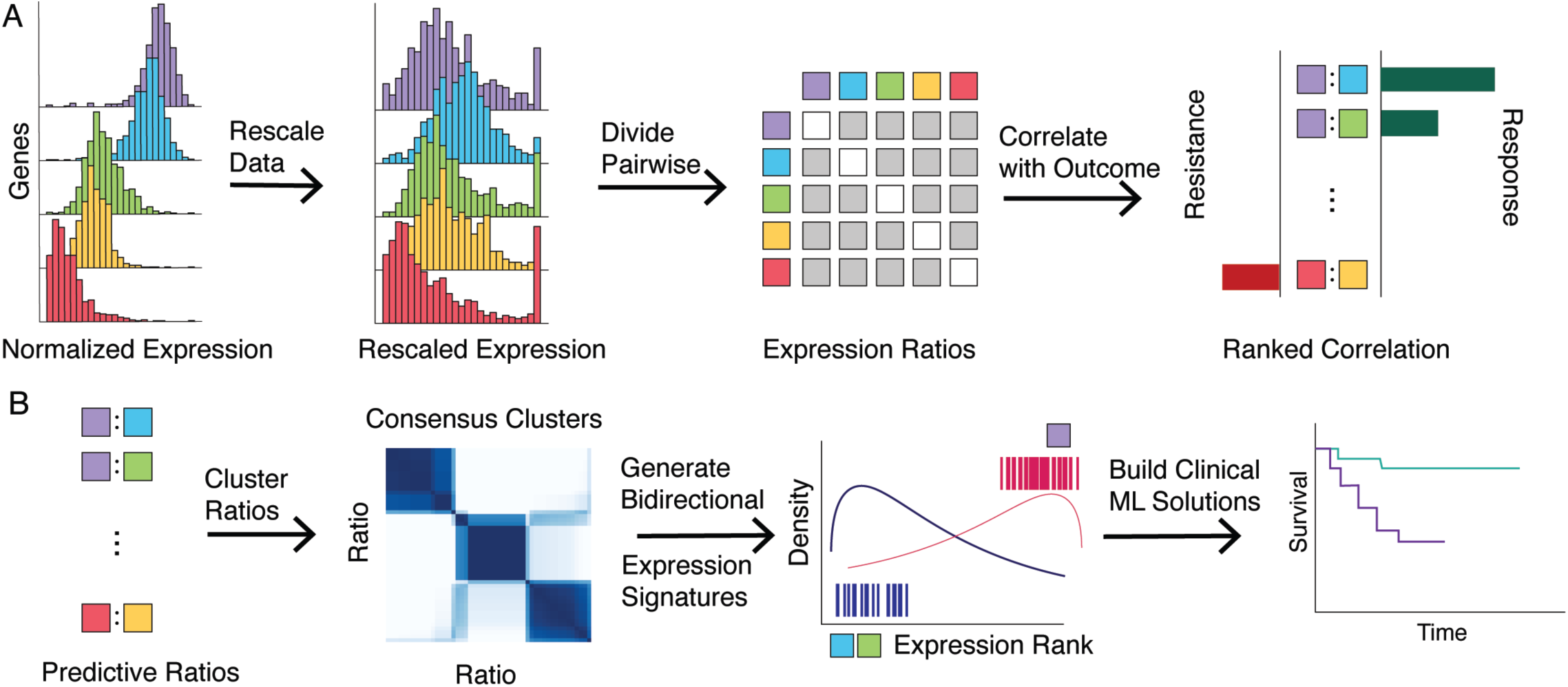
The tauX Approach for Generating Numerically Stable Gene Expression Ratios as Features for Clinical plications. First, gene ratios were calculated and associated with response to therapy (A). Gene ion counts were normalized to account for library size and transcript length (TPM). The TPM values were scaled to facilitate pairwise comparisons. The resulting feature matrix was then ranked to identify which re differentially expressed between responders and nonresponders. The response ratios were then ed to identify modules of ratio expression (B). The resulting gene ratio enrichment scores (GRES) were gineered into bidirectional gene sets and used as features for clinical ML applications.

Pairwise ratios were calculated across all genes of interest. Because the distribution of gene expression values varies considerably across genes, each gene was rescaled to a range between 1 and 100. Rescaling guarantees numerically stable comparisons across the entire transcriptome. Most pairwise comparisons were not associated with response, so a *t*-statistic was calculated to identify which ratios were consistently associated with response, accounting for the effect and sample sizes. Consensus clusters were optimized using recommended clustering quality metrics, including the BIC, the Davies–Bouldin index, the silhouette score, and the *Calinski–Harabasz* index (Monti et al., 2003). This resulted in the identification of 7 new response signatures for the StandUp2Cancer cohort (Supplementary Table 1). Each gene expression ratio module was further characterized using gene set enrichment analysis to characterize the functional enrichment of biological pathways. The gene ratio modules were then deconstructed into up- and downregulated gene sets for bidirectional gene set enrichment using the SingScore approach (Foroutan et al., 2018). The resulting enrichment scores greatly reduced the total number of features, which is beneficial, particularly when the amount of training data is small. The gene ratio enrichment scores (GRESs) were then used as features for training ML algorithms and applied to synthetic data and real-world lung cancer data.

### Performance of the tauX Approach on Synthetic Data

Synthetic datasets were constructed to evaluate the performance of the tauX approach compared to other widely used feature engineering strategies. These strategies were also compared across widely used machine learning algorithms, including ElasticNet, linear SVM, and nonlinear RBF SVM (Figure 2). Overall, the tauX strategy consistently outperformed the traditional approaches of using differentially expressed genes (DEGs) and highly variable genes (HVGs). The average AUCs for the tauX approach across the ElasticNet, linear SVM and RBF SVM models were 0.81, 0.83, and 0.83, respectively (Figure 2). The average AUC using the HVGs was 0.58 across all three methods. The average AUCs of the DEGs were 0.63, 0.61, and 0.61, respectively. The tauX signature was able to maintain predictive performance at lower effect sizes, suggesting that the tauX signature identifies subtle changes in expression that may correlate with the response to therapies.

**Figure 2.**
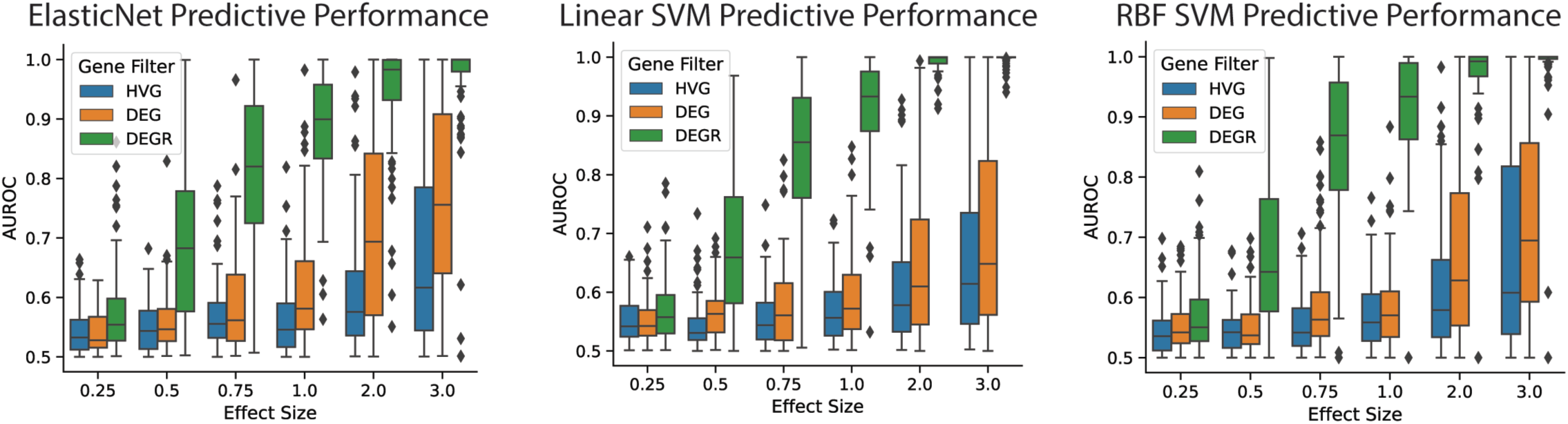
Average Predictive Performance using tauX Feature Engineering Strategy. The predictive performance, measured by area under the receiver operator curve (AUROC) for three feature engineering strategies: highly variable genes (HVG), differentially expressed genes (DEG) and differentially expressed gene ratios (DEGRs). The impact on predictive performance was assessed using the ElasticNet linear regression model, linear support vector machine, and a nonlinear RBF support vector machine.

Visual inspection of the features using hierarchically clustered heatmaps (Figure 3A) revealed considerable amounts of signals not associated with response for HVGs and DEGs. Surprisingly, the gene ratio approach showed low amounts of background signal and elevated levels of response-associated signal. The dendrogram for variably expressed genes and differentially expressed genes showed modest separation between responders and nonresponders, which is consistent with the observed clustering of other publicly available cancer drug response data (Hugo et al., 2016). The tauX-engineered features showed clear separation, suggesting that the ML algorithms may more easily identify these patterns (chi-squared p value < 0.05).

**Figure 3.**
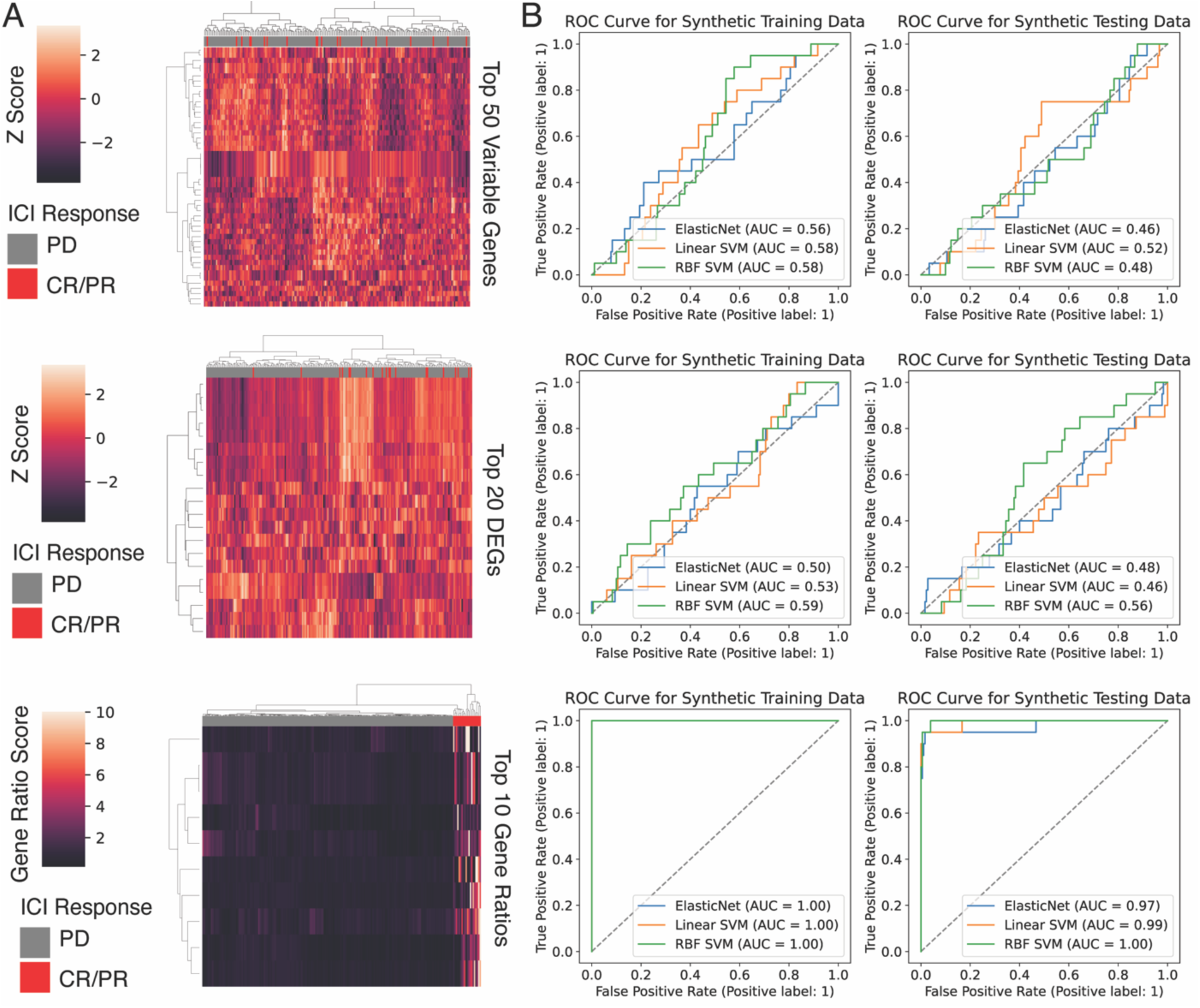
Visualization of Gene Expression Features and Resulting Predictive Performance. Heatmaps of engineered gene expression features for variably expressed (HVG), differentially expressed (DEG), and tauX transformed gene ratios (DEGR) (A). Resulting receiver operator curves (ROC) generated from training and testing the ElasticNet, linear SVM, and nonlinear SVM ML algorithms (B).

We next examined the predictive performance of commonly used ML algorithms. The ML algorithms were trained using Bayesian hyperparameter optimization on the training data and applied to a hold-out validation cohort (Figure 3B). The performance was visualized using a receiver operating characteristic (ROC) plot. The tauX engineered gene ratios achieved excellent performance on the training data (AUC > 0.9), whereas the traditional approaches were unable to identify a signal. To determine whether the tauX model overfit the training data, we predicted responses in an out-of-sample cohort of synthetic data generated from the same background cohort but not used for training. The tauX-generated features exhibited excellent predictive performance in the hold-out data (AUC > 0.9), suggesting that this strategy may have a computational advantage over existing gene expression feature engineering approaches.

### Statistical Properties Underlying the tauX Approach

We next investigated the statistical properties of the tauX strategy to explore how simple transformation of the gene expression data significantly improved its predictive performance. Synthetic data were generated to model a pair of inversely expressed genes, such as genes related to response and resistance marker expression (Supplementary Figure 3). One way to improve the specificity of an ML approach is to increase the distributional distance between responders and nonresponders. TauX transformation caused a significant increase in the distributional distance compared to the normalized expression distribution, as measured by the Kullback–Leibler divergence (Burnham & Anderson, 2001). Statistical modeling of the resulting ratio distribution revealed that the ratios follow an extreme value distribution instead of the original normal distribution, leading to significant differences between background and predictive response ratio expression. This transformation increases the expression differences between responders and nonresponders, which may facilitate more accurate and reproducible separation of responders by ML algorithms.

### Predicting ICI response in the stand-up-to-cancer (SU2C) lung cohort

The Stand Up To Cancer-Mark Foundation recently published an integrative analysis of non-small cell lung cancer (NSCLC) that included 152 samples with RNA-seq and immune checkpoint inhibitor (ICI) response data (Ravi et al., 2023). Preliminary exploration of established biomarkers of ICI response, including tumor mutation burden, neoantigen burden, and PDL1 expression, revealed moderate predictive performance (AUC < 0.8, Supplementary Figure 4). We sought to improve the predictive performance in this cohort by training ML models using tauX-generated GRES features. As a non-tauX-derived gene set comparator, we also assessed its predictive performance relative to the widely used hallmarks of cancer gene signatures (Liberzon et al., 2011).

Unsupervised consensus clustering of the responder gene expression cohort revealed three clear subgroups of patient samples (Supplementary Figure 2). An investigation of the histology of the clusters revealed that two of the clusters (specifically, clusters 0 and 2) correlated with LUAD histology, whereas cluster 1 was enriched for LUSC (chi-squared test, p value = 0.028). Upon further characterization of the responder clusters, it was found that cluster 0 samples were more likely to have a greater number of previous lines of therapy, whereas cluster 2 samples were more likely to have fewer previous lines of therapy (Mann‒ Whitney U test, p value = 0.002). The responder cluster assignments were used to randomly stratify the samples into training and validation datasets. This ensured that the training and validation cohorts had similar compositions of known gene expression covariates, including histological subtype, treatment history, and response to checkpoint blockade therapy.

The tauX approach was then applied to the training cohort (n=57). This resulted in the identification of 7 GRESs (Supplementary Table 1). An investigation of these GRESs revealed functional enrichment of biological pathways in the numerator and denominator gene sets (Figure 4). The LUAD patients with fewer previous lines of therapy (response cluster 2) were enriched for classical immune activation-associated gene sets in the numerator. Similar signatures have been shown to predict the response to checkpoint blockade therapy (Graham et al., 2021). However, not all patients with inflamed tumors respond to checkpoint blockade, so concomitant downregulation of resistance pathways may be required to achieve a response. The tauX approach also identified paired signatures of nonresponses, including transcriptional regulation, DNA damage repair, and lipid metabolism.

**Figure 4.**
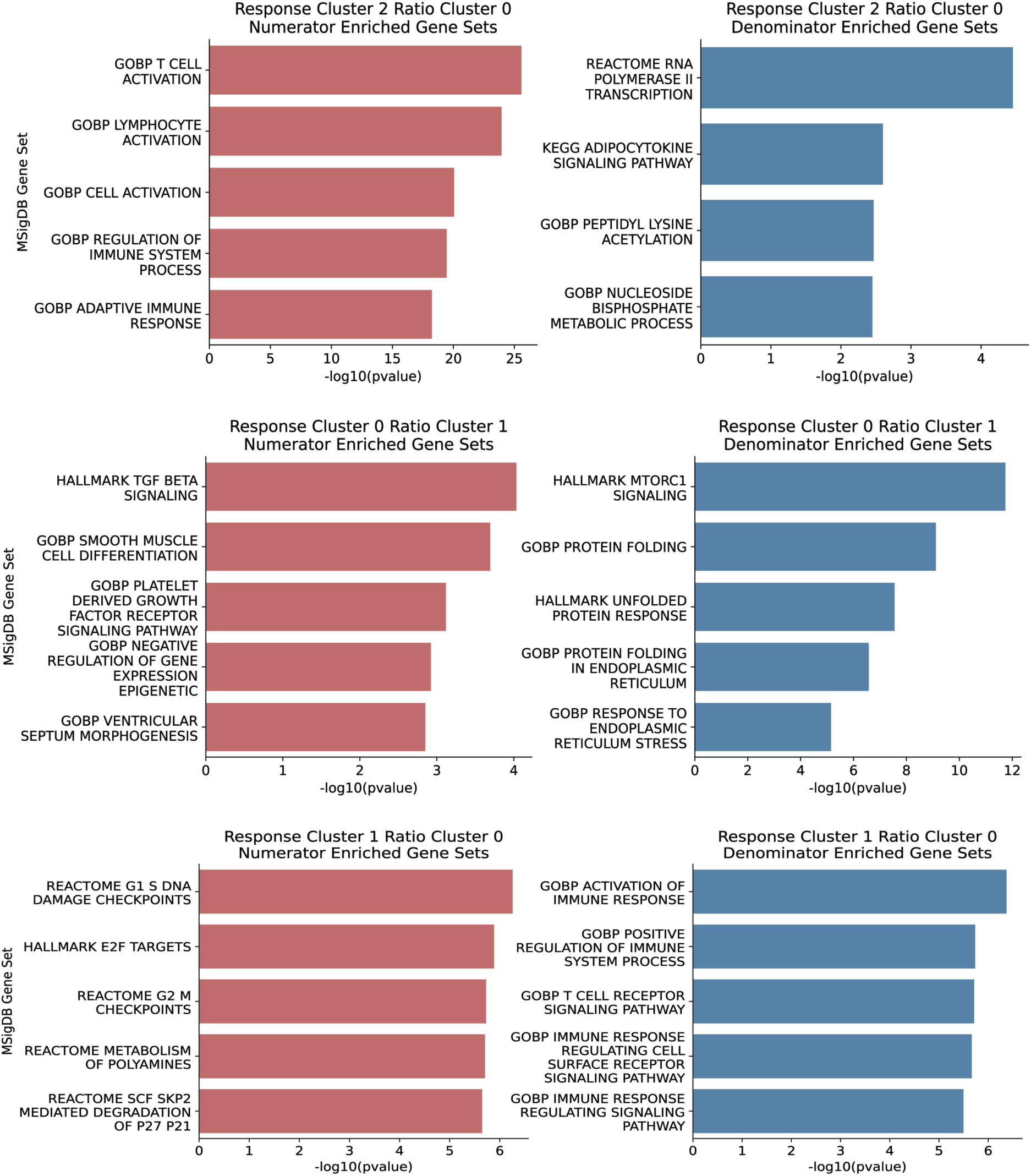
TauX Approach Applied to Stand Up 2 Cancer Lung ICI Dataset. Gene expression ratio values were clustered to find modules of ratio associated expression. Gene set enrichment analysis found coordinated expression of biological functions. The response clusters 0, 1, and 2 corresponded to LUAD/more treatments, LUSC, and LUAD/fewer treatments subgroups, respectively.

**Figure 5.**
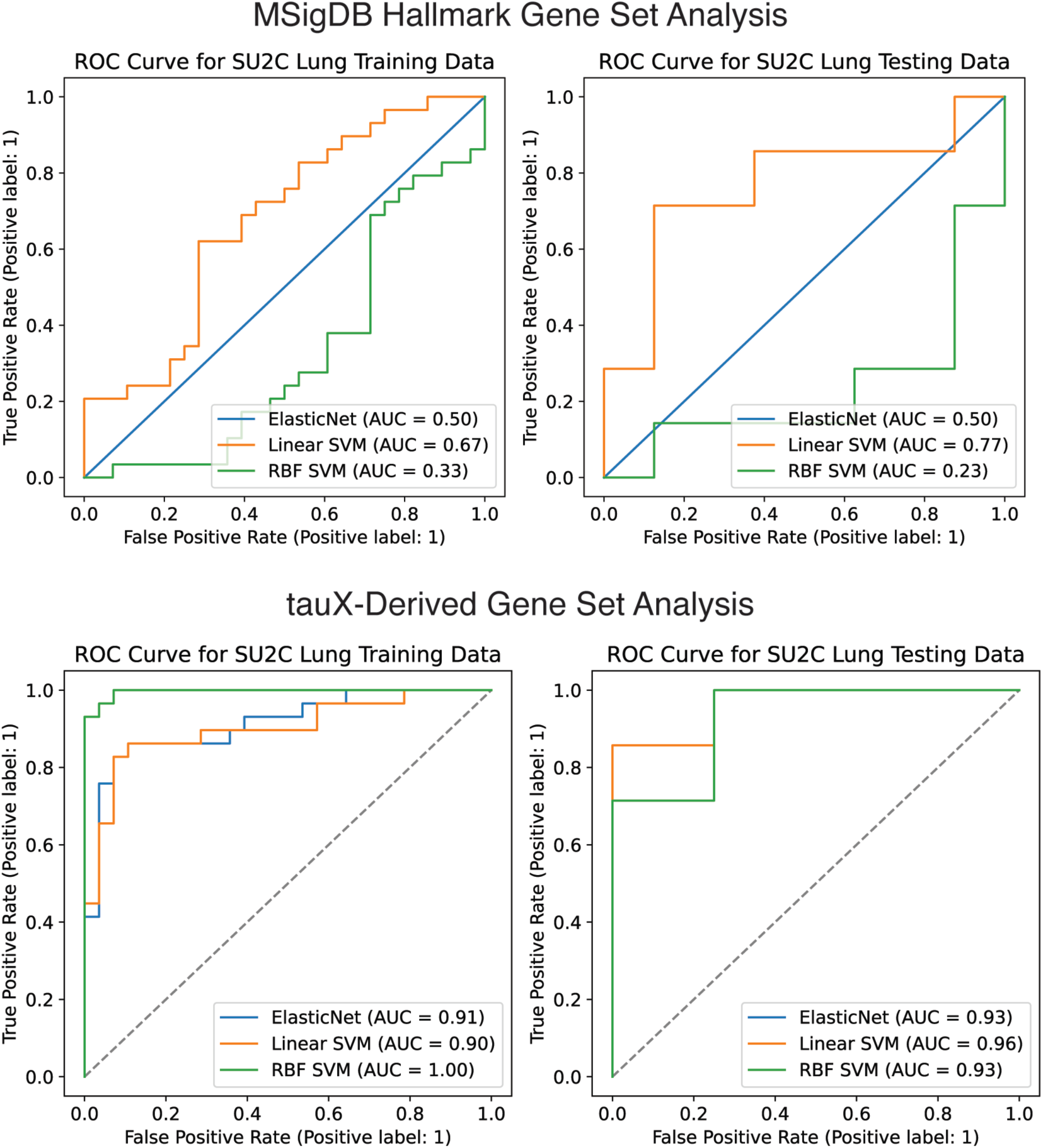
Predictive Performance Comparison between MSigDB Hallmark Gene Sets and the tauX-derived Gene Sets. Resulting gene set enrichment scores for the hallmark gene sets and the tauX gene sets were used for training ML algorithms. The predictive performance was visualized using the receiver operating characteristic (ROC) curves.

We found substantial enrichment in the more heavily pretreated LUAD patients for TGF-beta signaling, platelet growth factor receptor signaling, smooth muscle differentiation, and endoplasmic reticulum transport signaling (Figure 4). The cluster 0 LUAD denominator gene set was enriched for protein folding pathways and endoplasmic reticulum stress pathways, which have been shown to be involved in resistance to cancer therapy through adaptation to hypoxia, inflammation, and angiogenesis (Yadav et al., 2014). Finally, the LUSC cluster showed the inverse trend to that of the LUAD clusters, where cell cycle signaling was a positive predictor of response and immune cell expression was a predictor of resistance. A similar pattern has also been observed by others as a resistance signature in LUSC (Yang et al., 2022).

As a comparator to the tauX-defined GRESs, we evaluated performance using the cancer hallmark gene set signatures as input to the same set of ML algorithms used above (Liberzon et al., 2011). Hallmark gene sets are widely used to characterize cancer gene expression, as these gene sets were defined to capture important expression patterns associated with cancer biology and the tumor microenvironment (Memon et al., 2024). Previously, hallmark models were built using Bayesian hyperparameter optimization to determine the best performing models for each of the feature sets. The hallmark linear SVM achieved an AUC comparable to that of existing biomarkers of response, including PDL1 staining and the TMB (AUC ∼ 0.7, Supplementary Figure 4). The models trained using enrichment scores for the 7 previously defined tauX features achieved superior predictive performance for the training and validation cohorts (AUC > 0.9), suggesting that the tauX approach may isolate more informative features for training clinical ML tools in the SU2C anti-PD1 response cohort.

### The tauX signature was correlated with patient survival in the SU2C and TCGA LUAD cohorts

The tauX approach for this lung cancer cohort was initially trained on a binary outcome variable associated with response. To evaluate patient survival, which is not necessarily equivalent to response, we assessed whether the tauX predictions also correlated with greater progression-free survival in the SU2C cohort. A significant difference in survival outcomes was observed in the validation cohort for progression-free survival using the elastic net and linear SVM classifiers (log-rank test, *p* value = 0.0012 and 0.0062, respectively) (Figure 6) but not the RBF SVM classifier, although the survival curves showed a similar pattern (log-rank test, p value = 0.12). We then investigated whether tauX GRESs also predict patient survival in The Cancer Genome Atlas (TCGA-LUAD) LUAD cohort (Collisson et al., 2014). A Cox proportional hazards model was fit to the TCGA-LUAD cohort using the corresponding LUAD tauX signatures and the nonsilent mutation rate as covariates (Figure 6). The response cluster 2 signature was the most relevant signature for the treatment-naïve TCGA-LUAD cohort and achieved the greatest reduction in relative risk (HR 95% CI: 0.13-0.64). Interestingly, the nonsilent mutation rate was not an independent covariate for decreased risk when the response cluster 2 signature was included as a covariate (HR 95% CI: 0.97-1.01).

**Figure 6.**
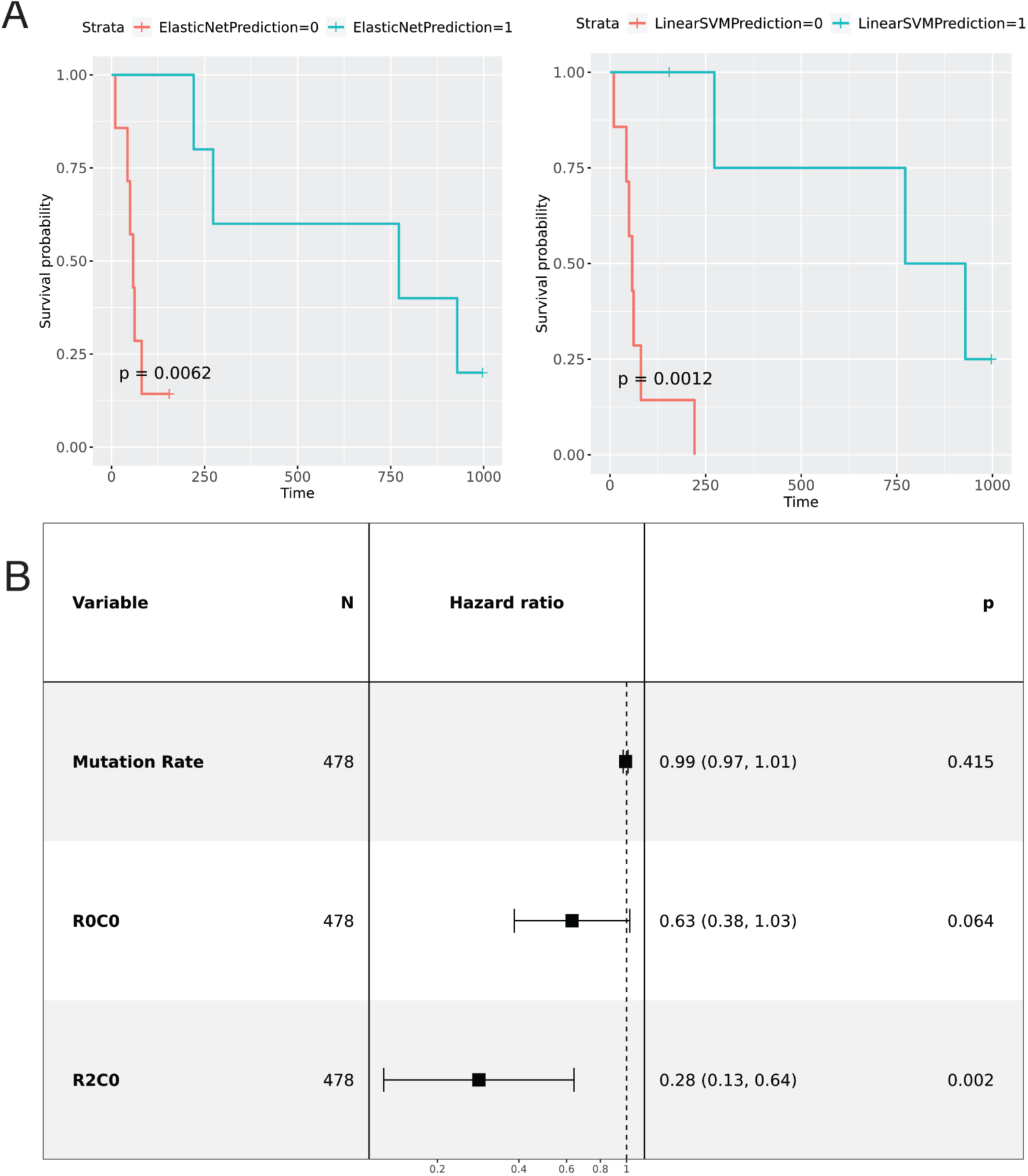
Survival Analysis using the tauX ICI Response Signatures. Kaplan‒Meier plots for the SU2C Lung validation cohort showing the difference in survival curves for tauX predicted responders and nonresponders (A). Forest plot of the Cox proportional hazards model parameters for the TCGA LUAD survival data comparing nonsilent mutation rate, the response subtype 0 cluster 0 tauX signature (R0C0), and the response subtype 2 cluster 0 tauX signature (R2C0) as covariates (B).

## Discussion

Gene expression analysis paired with ML tools has transformative potential for patient care but requires substantial optimization to overcome well-documented issues with overfitting and reproducibility. One strategy for ML tool performance optimization, which is based on the recognition that both the response to therapy and gene expression regulatory structure are affected by positive and negative factors, may be related to putatively relevant gene expression signals (Sinicrope et al., 2020; Turan et al., 2021). Here, we showed that a transformation of gene expression counts into ratios has the potential to amplify subtle changes in expression that correlate with the response to ICI therapy. To our knowledge, this is the first evaluation of pairwise gene expression ratios for ML tool optimization in the context of response prediction. Using these ratios, we identified correlated modules associated with positive and negative predictive factors for ICI response and showed that these gene ratio expression signatures (GRESs) can be used to enhance the predictive performance of ML tools relative to the commonly used hallmarks of cancer gene set collection.

The tauX strategy is built upon a simple idea that has deep biological significance. Gene expression is highly correlated with the coregulation of many genes and pathways to control gene activity. When one gene or pathway is activated, the anti-gene/pathway pathway is deactivated (i.e., Ras GEFs vs Ras GAPs). This pattern can also be seen in the response prediction data, where the response markers are correlated with the nonresponse markers. This makes prediction challenging when a positive response prediction score may be cancelled out by an increase in the resistance prediction score. The tauX approach models the response- and resistance-associated expression simultaneously to improve its predictive accuracy. Overexpressed and underexpressed genes are regularly characterized during routine differential expression analysis. Combining positive and negative predictors via pairwise comparisons may be a new way to generate clinical gene expression signatures.

The SU2C lung cohort included patients with diverse treatment histories, and we were able to leverage this advantage here to isolate a distinct response signature for more heavily treated patients. This is a particularly exciting discovery since heavily pretreated populations of patients tend to be less responsive to checkpoint blockade therapy (Huang et al., 2023). The tauX approach revealed elevated TGF-beta signaling, with concurrent downregulation of endoplasmic reticulum stress response pathways being associated with the response. The ratios driving TGF-beta enrichment included those of the SKIL gene, which is upregulated during sustained TGF-beta signaling (Tecalco-Cruz et al., 2012). TGF-beta signaling is currently considered a predictor of nonresponse to checkpoint blockade therapy and predicts worse survival in patients with lung cancer (Stefanescu et al., 2021; Tschernia & Gulley, 2022). TGF-beta signaling may maintain an exhausted T-cell state through the inhibition of stem cell-like CD8+ T cells. Simultaneous PDL1 and TGF-beta blockade was shown to overcome these resistance mechanisms and allow antitumor immune responses to eradicate tumors (Castiglioni et al., 2023). The unfolded protein response (UPR) has been shown to make tumors more resilient to cellular stress in the tumor microenvironment and facilitate immune evasion through signaling to tumor-promoting monocytes/macrophages (Zanetti et al., 2022). The tauX approach provides more context to the response signature and provides insight into mechanisms that are not as easily identified using traditional gene expression analysis.

The largest cluster of patients in the SU2C cohort consisted of LUAD patients with fewer previous lines of therapy (<= 2 lines). This cluster was associated with classic predictors of response to checkpoint blockade, including T-cell activation signatures (X. Li et al., 2019). The tauX approach enhanced the adaptive immune response signature by pairing it with gene expression patterns associated with resistance, which included upregulation of proliferative signatures (Cristescu et al., 2018). This cluster clinically most closely resembled the pretreatment naïve TCGA LUAD cohort, and indeed, the tauX predictions were associated with better survival outcomes. This finding is consistent with the known observation that the immune involvement associated with checkpoint blockade response also correlates with a survival benefit in pretreated TCGA samples (Thorsson et al., 2018).

Many methods have been developed to address heterogeneity in RNA-seq data, but few of these methods are specifically designed for clinical applications. The unique constraints of medical gene expression analysis require a new set of tools to achieve clinical impact. We designed tauX to be applied to any ML task that has a binary outcome variable. This approach may be particularly helpful for relatively small training cohorts (< 100 samples), since tauX transformation improves specificity by increasing the distributional distance between responders and nonresponders. The tauX strategy leverages the statistical properties of gene expression data to learn new gene expression signatures associated with response.

In conclusion, we present the tauX approach as a flexible modeling strategy with the potential to improve the performance of ML tools for response prediction using gene expression data. We have published the tauX approach as a docker container to enable further community development of this framework, with the hope that this approach will contribute to the advancement of ML for precision medicine applications.

## Declarations

### Availability of data and materials

All the data and Jupyter notebooks used to generate the results for the manuscript are available for review at a publicly available GitHub repository (https://github.com/jpfeil/taux-manuscript). The tauX feature generator is available as a stable docker container that can be pulled from dockerhub (https://hub.docker.com/r/pfeiljx/taux). The TCGA and SU2C data are publicly available.

## AbbVie Disclosure Statement

All authors are current employees of AbbVie. The design, study design, and financial support for this research were provided by AbbVie. AbbVie participated in the interpretation of the data and the review and approval of the publication.

## Supplemental Figures

**Supplementary Figure 1.**
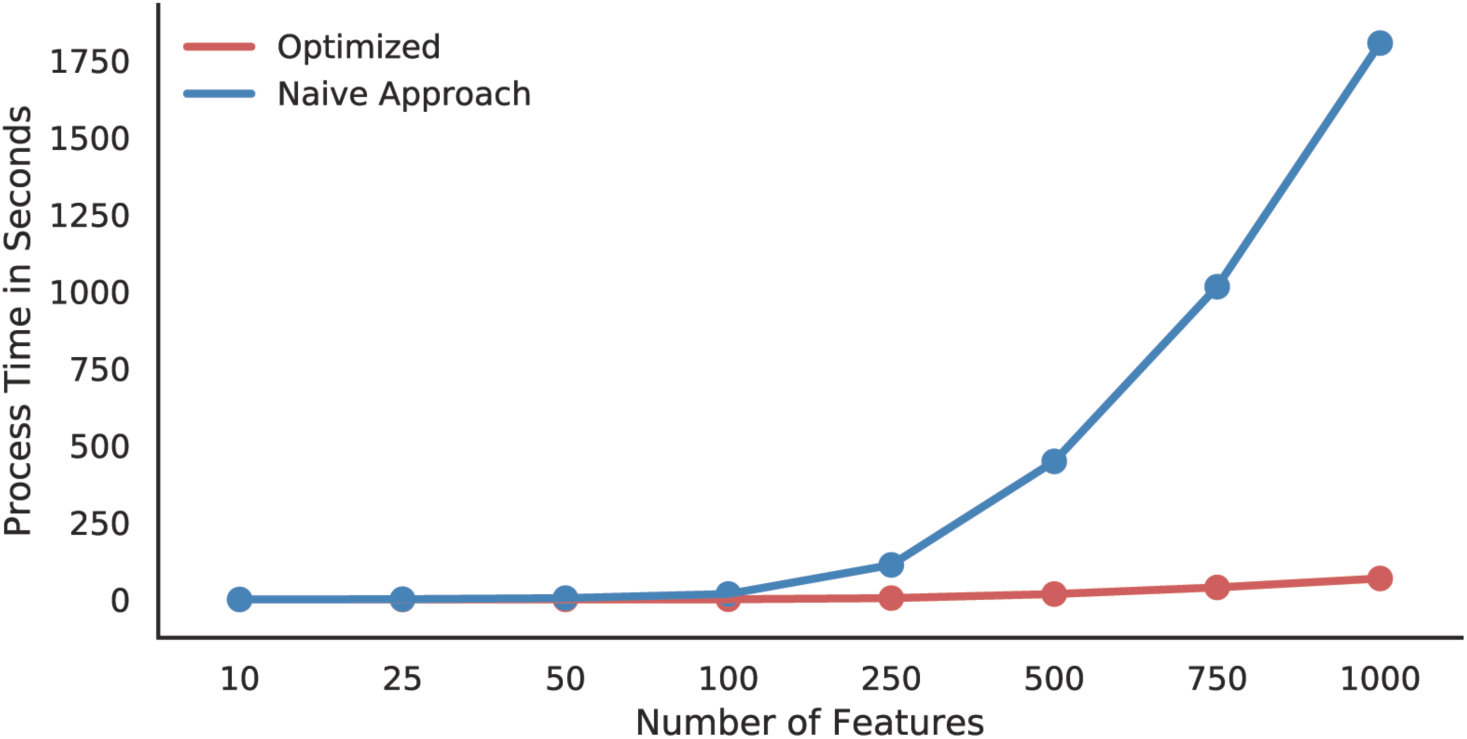
Performance improvement over naïve ratio calculation allows for more comprehensive search for novel expression markers. Synthetic gene expression data for a cohort of 100 samples were created for a range of feature sizes, ranging from 10 to 1000 features. The optimized matrix algebra approach scaled linearly with the number of features, whereas the time needed to achieve the naive approach grew exponentially even for a relatively small number of genes.

**Supplementary Figure 2.**
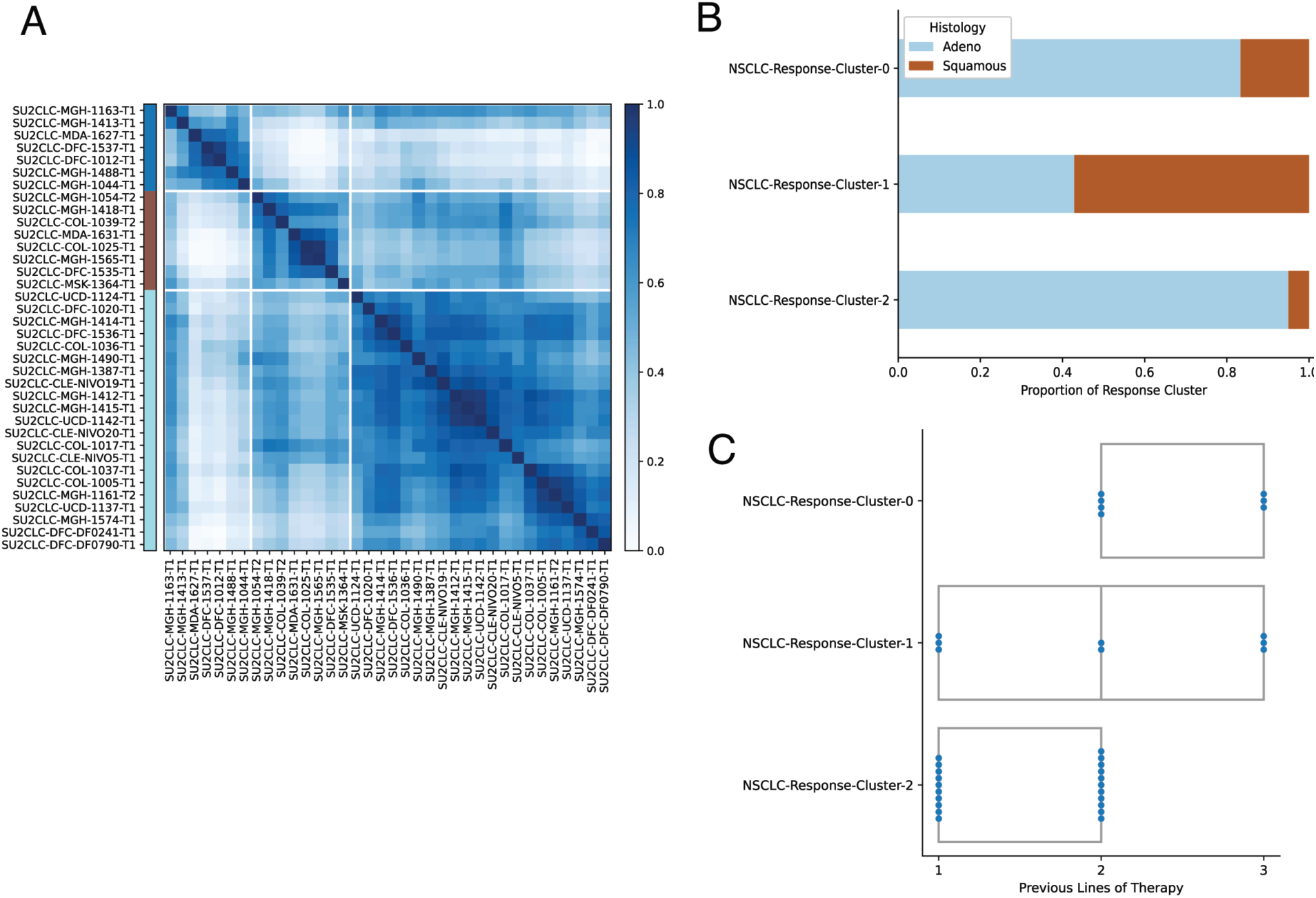
Significant clinical differences between SU2C response clusters. Checkpoint blockade responders were consensus clustered (A). Stacked barplot of histology annotations for responder clusters (B). Responder clusters were statistically associated with lung histology (Chi-squared p value < 0.05). Boxplots for number of lines of therapy for each response cluster (C). Numbers of lines of therapy were significantly different between the LUAD cluster 0 and cluster 2 (Mann‒Whitney U test p value < 0.05).

**Supplementary Figure 3.**
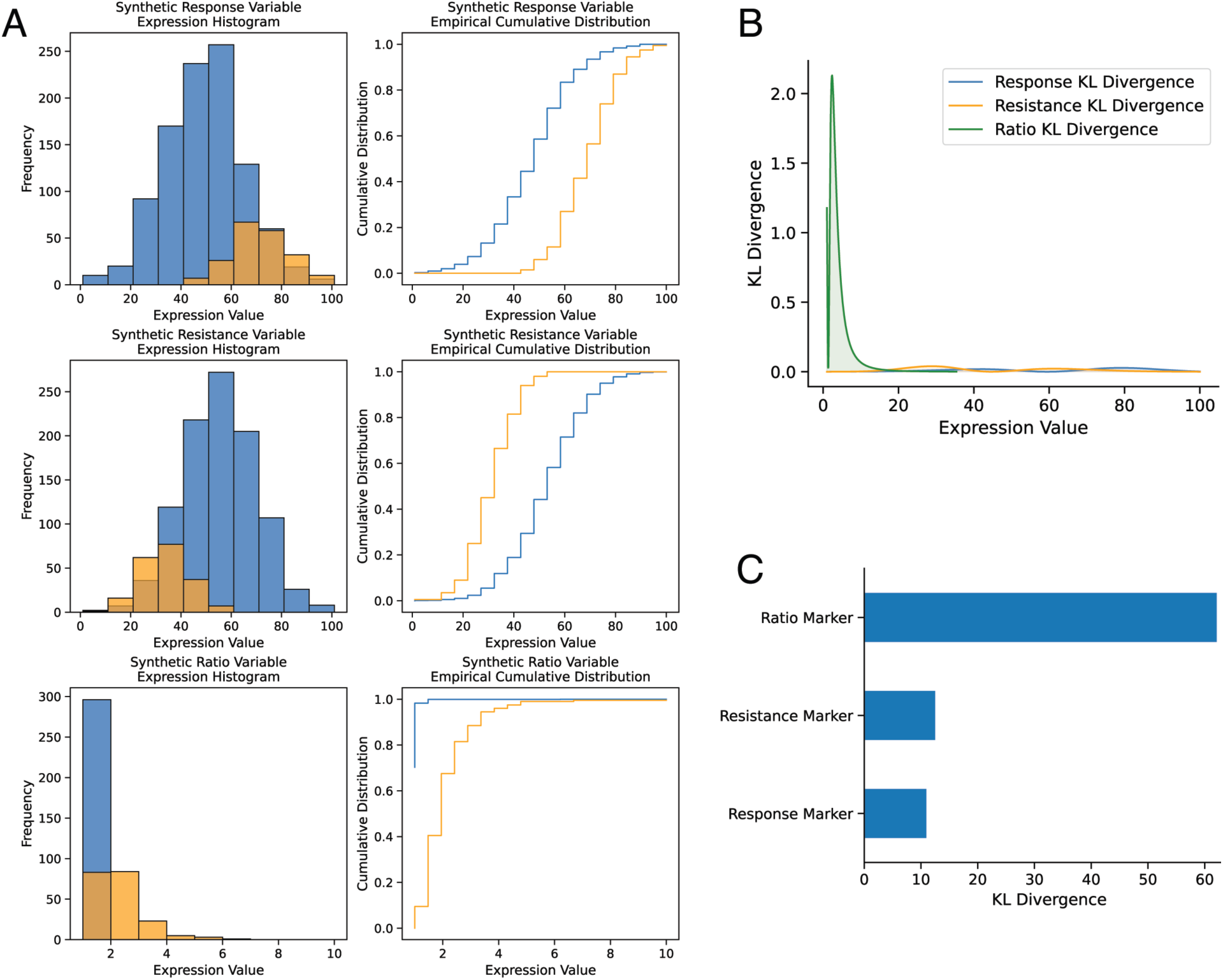
Statistical Analysis of tauX Approach. Histogram and cumulative distribution plots for synthetic “responder” and “nonresponder” data (A). Kullback‒Leibler divergence values across expression values (B). Summed Kullback‒Leibler divergence barplots as a measure of distributional distance across features (C).

**Supplementary Figure 4.**
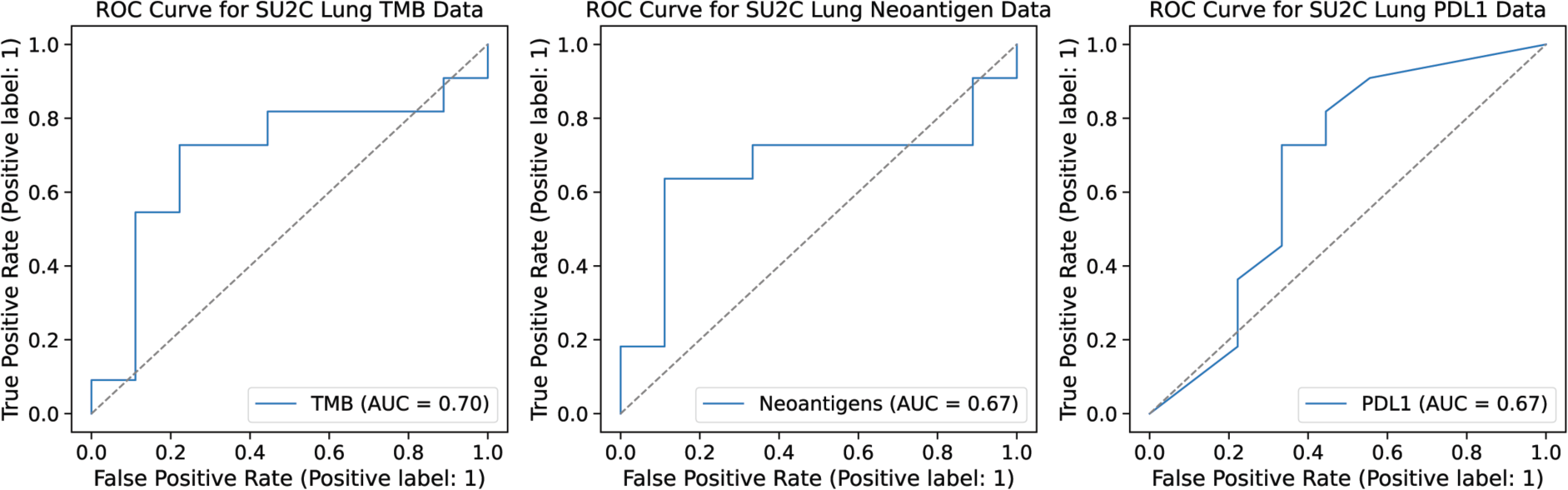
SU2C ROC Plots for Existing Checkpoint Blockade Response Markers. Receiver operating characteristics (ROC) plots for existing biomarkers of response to checkpoint blockade therapy.

**Supplementary Figure 5.**
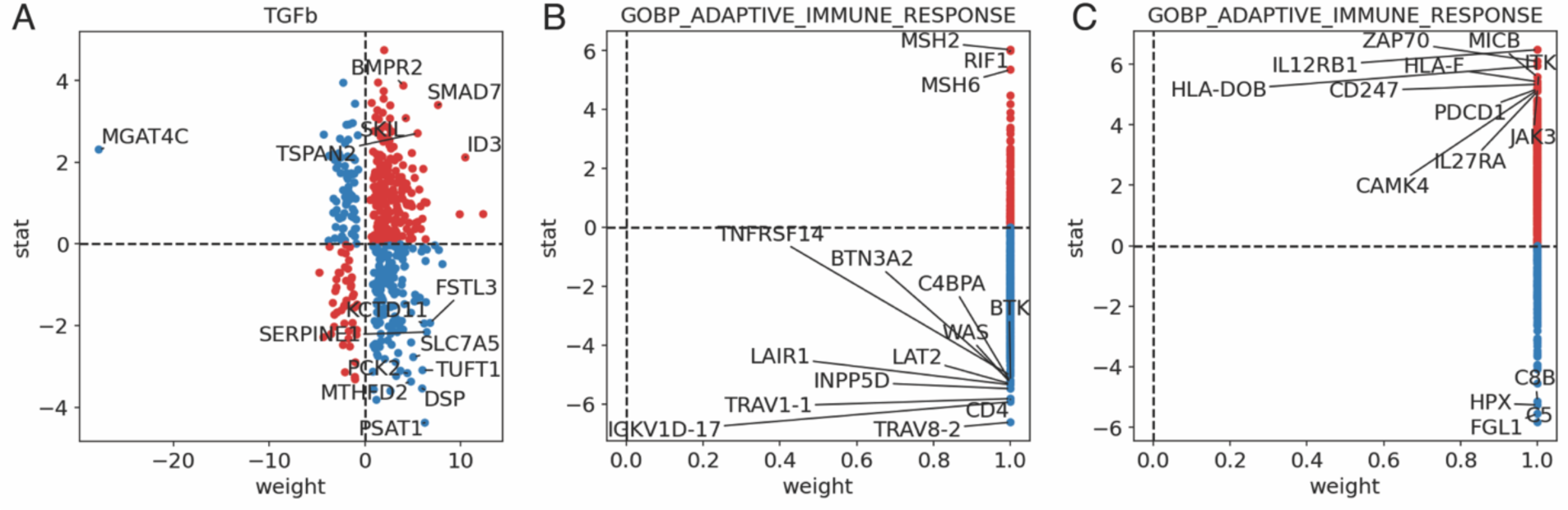
Gene Expression Target Plots for SU2C Lung Response Clusters. Progeny TGFb gene set expression for the SU2C lung response cluster 0. GO biological process gene set expression for SU2C lung response cluster 1. GO biological process gene set expression for SU2C lung response cluster 2.

**Supplementary Table 1** Supplementary-table-1.xlsx

## References

Akiba, T., Sano, S., Yanase, T., Ohta, T., & Koyama, M. (2019). Optuna: A Next-generation Hyperparameter Optimization Framework. Proceedings of the 25th ACM SIGKDD International Conference on Knowledge Discovery & Data Mining, 2623–2631. 10.1145/3292500.3330701

Altman, N., & Krzywinski, M. (2018). The curse(s) of dimensionality. Nature Methods, 15(6), 399–400. 10.1038/s41592-018-0019-x

Anders, S., & Huber, W. (2010). Differential expression analysis for sequence count data. Genome Biology, 11(10), R106. 10.1186/gb-2010-11-10-r106

Ashburner, M., Ball, C. A., Blake, J. A., Botstein, D., Butler, H., Cherry, J. M., Davis, A. P., Dolinski, K., Dwight, S. S., Eppig, J. T., Harris, M. A., Hill, D. P., Issel-Tarver, L., Kasarskis, A., Lewis, S., Matese, J. C., Richardson, J. E., Ringwald, M., Rubin, G. M., & Sherlock, G. (2000). Gene Ontology: Tool for the unification of biology. Nature Genetics, 25(1), Article 1. 10.1038/75556

Badia-i-Mompel, P., Vélez Santiago, J., Braunger, J., Geiss, C., Dimitrov, D., Müller-Dott, S., Taus, P., Dugourd, A., Holland, C. H., Ramirez Flores, R. O., & Saez-Rodriguez, J. (2022). decoupleR: Ensemble of computational methods to infer biological activities from omics data. Bioinformatics Advances, 2(1), vbac016. 10.1093/bioadv/vbac016

Barbie, D. A., Tamayo, P., Boehm, J. S., Kim, S. Y., Moody, S. E., Dunn, I. F., Schinzel, A. C., Sandy, P., Meylan, E., Scholl, C., Fröhling, S., Chan, E. M., Sos, M. L., Michel, K., Mermel, C., Silver, S. J., Weir, B. A., Reiling, J. H., Sheng, Q., … Hahn, W. C. (2009). Systematic RNA interference reveals that oncogenic KRAS-driven cancers require TBK1. Nature, 462(7269), Article 7269. 10.1038/nature08460

Bar-Joseph, Z., Gifford, D. K., & Jaakkola, T. S. (2001). Fast optimal leaf ordering for hierarchical clustering. Bioinformatics, 17(suppl_1), S22–S29. 10.1093/bioinformatics/17.suppl_1.S22

Bretthauer, M., Wieszczy, P., Løberg, M., Kaminski, M. F., Werner, T. F., Helsingen, L. M., Mori, Y., Holme, Ø., Adami, H.-O., & Kalager, M. (2023). Estimated Lifetime Gained With Cancer Screening Tests: A Meta-Analysis of Randomized Clinical Trials. JAMA Internal Medicine. 10.1001/jamainternmed.2023.3798

Breunig, M. M., Kriegel, H.-P., Ng, R. T., & Sander, J. (2000). LOF: Identifying density-based local outliers. Proceedings of the 2000 ACM SIGMOD International Conference on Management of Data, 93–104.

Burnham, K. P., & Anderson, D. R. (2001). Kullback‒Leibler information as a basis for strong inference in ecological studies. Wildlife Research, 28(2), 111–119. 10.1071/wr99107

Campbell, J. D., Alexandrov, A., Kim, J., Wala, J., Berger, A. H., Pedamallu, C. S., Shukla, S. A., Guo, G., Brooks, A. N., Murray, B. A., Imielinski, M., Hu, X., Ling, S., Akbani, R., Rosenberg, M., Cibulskis, C., Ramachandran, A., Collisson, E. A., Kwiatkowski, D. J., … Meyerson, M. (2016). Distinct patterns of somatic genome alterations in lung adenocarcinomas and squamous cell carcinomas. Nature Genetics, 48(6), Article 6. 10.1038/ng.3564

Castiglioni, A., Yang, Y., Williams, K., Gogineni, A., Lane, R. S., Wang, A. W., Shyer, J. A., Zhang, Z., Mittman, S., Gutierrez, A., Astarita, J. L., Thai, M., Hung, J., Yang, Y. A., Pourmohamad, T., Himmels, P., De Simone, M., Elstrott, J., Capietto, A.-H., … Müller, S. (2023). Combined PD-L1/TGFβ blockade allows expansion and differentiation of stem cell-like CD8 T cells in immune excluded tumors. Nature Communications, 14(1), Article 1. 10.1038/s41467-023-40398-4

Cheadle, C., Vawter, M. P., Freed, W. J., & Becker, K. G. (2003). Analysis of Microarray Data Using Z Score Transformation. The Journal of Molecular Diagnostics : JMD, 5(2), 73–81.

Collisson, E. A., Campbell, J. D., Brooks, A. N., Berger, A. H., Lee, W., Chmielecki, J., Beer, D. G., Cope, L., Creighton, C. J., Danilova, L., Ding, L., Getz, G., Hammerman, P. S., Neil Hayes, D., Hernandez, B., Herman, J. G., Heymach, J. V., Jurisica, I., Kucherlapati, R., … John Flynn Hospital. (2014). Comprehensive molecular profiling of lung adenocarcinoma. Nature, 511(7511), Article 7511. 10.1038/nature13385

Cristescu, R., Mogg, R., Ayers, M., Albright, A., Murphy, E., Yearley, J., Sher, X., Liu, X. Q., Lu, H., Nebozhyn, M., Zhang, C., Lunceford, J. K., Joe, A., Cheng, J., Webber, A. L., Ibrahim, N., Plimack, E. R., Ott, P. A., Seiwert, T. Y., … Kaufman, D. (2018). Pan-tumor genomic biomarkers for PD-1 checkpoint blockade–based immunotherapy. Science, 362(6411), eaar3593. 10.1126/science.aar3593

Foroutan, M., Bhuva, D. D., Lyu, R., Horan, K., Cursons, J., & Davis, M. J. (2018). Single sample scoring of molecular phenotypes. BMC Bioinformatics, 19(1), 404. 10.1186/s12859-018-2435-4

Gillespie, M., Jassal, B., Stephan, R., Milacic, M., Rothfels, K., Senff-Ribeiro, A., Griss, J., Sevilla, C., Matthews, L., Gong, C., Deng, C., Varusai, T., Ragueneau, E., Haider, Y., May, B., Shamovsky, V., Weiser, J., Brunson, T., Sanati, N., … D’Eustachio, P. (2022). The reactome pathway knowledgebase 2022. Nucleic Acids Research, 50(D1), D687–D692. 10.1093/nar/gkab1028

Graham, L. S., Pritchard, C. C., & Schweizer, M. T. (2021). Hypermutation, Mismatch Repair Deficiency, and Defining Predictors of Response to Checkpoint Blockade. Clinical Cancer Research, 27(24), 6662–6665. 10.1158/1078-0432.CCR-21-3031

Hänzelmann, S., Castelo, R., & Guinney, J. (2013). GSVA: Gene set variation analysis for microarray and RNA-Seq data. BMC Bioinformatics, 14(1), 7. 10.1186/1471-2105-14-7

Huang, Y., Zhao, J. J., Soon, Y. Y., Kee, A., Tay, S. H., Aminkeng, F., Ang, Y., Wong, A. S. C., Bharwani, L. D., Goh, B. C., & Soo, R. A. (2023). Factors Predictive of Primary Resistance to Immune Checkpoint Inhibitors in Patients with Advanced Non-Small Cell Lung Cancer. Cancers, 15(10), 2733. 10.3390/cancers15102733

Hugo, W., Zaretsky, J. M., Sun, L., Song, C., Moreno, B. H., Hu-Lieskovan, S., Berent-Maoz, B., Pang, J., Chmielowski, B., Cherry, G., Seja, E., Lomeli, S., Kong, X., Kelley, M. C., Sosman, J. A., Johnson, D. B., Ribas, A., & Lo, R. S. (2016). Genomic and Transcriptomic Features of Response to Anti-PD-1 Therapy in Metastatic Melanoma. Cell, 165(1), 35–44. 10.1016/j.cell.2016.02.065

Hunter, J. D. (2007). Matplotlib: A 2D graphics environment. Computing in Science & Engineering, 9(3), 90–95. 10.1109/MCSE.2007.55

Kanehisa, M., & Goto, S. (2000). KEGG: Kyoto Encyclopedia of Genes and Genomes. Nucleic Acids Research, 28(1), 27–30. 10.1093/nar/28.1.27

Kassambara, A., Kosinski, M., Biecek, P., & Fabian, S. (2021). survminer: Drawing Survival Curves using “ggplot2” (0.4.9) [Computer software]. https://cran.r-project.org/web/packages/survminer/index.html

Lai, Y., He, S., Lin, Z., Yang, F., Zhou, Q., & Zhou, X. (2021). An Adaptive Robust Semi-Supervised Clustering Framework Using Weighted Consensus of Random kk-Means Ensemble. IEEE Transactions on Knowledge and Data Engineering, 33(5), 1877–1890. 10.1109/TKDE.2019.2952596

Li, K.-L., Huang, H.-K., Tian, S.-F., & Xu, W. (2003). Improving one-class SVM for anomaly detection. Proceedings of the 2003 International Conference on Machine Learning and Cybernetics (IEEE Cat. No.03EX693), 5, 3077–3081 Vol.5. 10.1109/ICMLC.2003.1260106

Li, X., Song, W., Shao, C., Shi, Y., & Han, W. (2019). Emerging predictors of the response to the blockade of immune checkpoints in cancer therapy. Cellular and Molecular Immunology, 16(1), 28–39. 10.1038/s41423-018-0086-z

Liberzon, A., Subramanian, A., Pinchback, R., Thorvaldsdóttir, H., Tamayo, P., & Mesirov, J. P. (2011). Molecular signatures database (MSigDB) 3.0. Bioinformatics, 27(12), 1739–1740. 10.1093/bioinformatics/btr260

Liu, F. T., Ting, K. M., & Zhou, Z.-H. (2008). Isolation Forest. 2008 Eighth IEEE International Conference on Data Mining, 413–422. 10.1109/ICDM.2008.17

Lun, A. T. L., McCarthy, D. J., & Marioni, J. C. (2016). A step-by-step workflow for low-level analysis of single-cell RNA-seq data with Bioconductor. F1000Research, 5, 2122. 10.12688/f1000research.9501.2

Mason, M., Lapuente-Santana, Ó., Halkola, A. S., Wang, W., Mall, R., Xiao, X., Kaufman, J., Fu, J., Pfeil, J., Banerjee, J., Chung, V., Chang, H., Chasalow, S. D., Lin, H. Y., Chai, R., Yu, T., Finotello, F., Mirtti, T., Mäyränpää, M. I., … Vincent, B. G. (2022). A Community Challenge to Predict Clinical Outcomes After Immune Checkpoint Blockade in Non-Small Cell Lung Cancer (p. 2022.12.05.518667). bioRxiv. 10.1101/2022.12.05.518667

Memon, D., Schoenfeld, A. J., Ye, D., Fromm, G., Rizvi, H., Zhang, X., Keddar, M. R., Mathew, D., Yoo, K. J., Qiu, J., Lihm, J., Miriyala, J., Sauter, J. L., Luo, J., Chow, A., Bhanot, U. K., McCarthy, C., Vanderbilt, C. M., Liu, C., … Hellmann, M. D. (2024). Clinical and molecular features of acquired resistance to immunotherapy in non-small cell lung cancer. *Cancer Cell*, S1535610823004415. 10.1016/j.ccell.2023.12.013

Monti, S., Tamayo, P., Mesirov, J., & Golub, T. (2003). Consensus Clustering: A Resampling-Based Method for Class Discovery and Visualization of Gene Expression Microarray Data. Machine Learning, 52(1), 91–118. 10.1023/A:1023949509487

Morozova, O., Hirst, M., & Marra, M. A. (2009). Applications of New Sequencing Technologies for Transcriptome Analysis. Annual Review of Genomics and Human Genetics, 10(1), 135–151. 10.1146/annurev-genom-082908-145957

Ott, T. (2022). *TankredO/pyckmeans:* [Computer software]. Zenodo. 10.5281/zenodo.6470080

Pedregosa, F., Varoquaux, G., Gramfort, A., Michel, V., Thirion, B., Grisel, O., Blondel, M., Prettenhofer, P., Weiss, R., Dubourg, V., Vanderplas, J., Passos, A., Cournapeau, D., Brucher, M., Perrot, M., & Duchesnay, É. (2011). Scikit-learn: Machine Learning in Python. Journal of Machine Learning Research, 12(85), 2825–2830.

Ravi, A., Hellmann, M. D., Arniella, M. B., Holton, M., Freeman, S. S., Naranbhai, V., Stewart, C., Leshchiner, I., Kim, J., Akiyama, Y., Griffin, A. T., Vokes, N. I., Sakhi, M., Kamesan, V., Rizvi, H., Ricciuti, B., Forde, P. M., Anagnostou, V., Riess, J. W., … Gainor, J. F. (2023). Genomic and transcriptomic analysis of checkpoint blockade response in advanced non-small cell lung cancer. Nature Genetics, 1–13. 10.1038/s41588-023-01355-5

Rodon, J., Soria, J.-C., Berger, R., Miller, W. H., Rubin, E., Kugel, A., Tsimberidou, A., Saintigny, P., Ackerstein, A., Braña, I., Loriot, Y., Afshar, M., Miller, V., Wunder, F., Bresson, C., Martini, J.-F., Raynaud, J., Mendelsohn, J., Batist, G., … Kurzrock, R. (2019). Genomic and transcriptomic profiling expands precision cancer medicine: The WINTHER trial. Nature Medicine, 25(5), 751–758. 10.1038/s41591-019-0424-4

Shah, P., Kendall, F., Khozin, S., Goosen, R., Hu, J., Laramie, J., Ringel, M., & Schork, N. (2019). Artificial intelligence and machine learning in clinical development: A translational perspective. Npj Digital Medicine, 2(1), Article 1. 10.1038/s41746-019-0148-3

Sinicrope, F. A., Shi, Q., Hermitte, F., Zemla, T. J., Mlecnik, B., Benson, A. B., Gill, S., Goldberg, R. M., Kahlenberg, M. S., Nair, S. G., Shields, A. F., Smyrk, T. C., Galon, J., & Alberts, S. R. (2020). Contribution of Immunoscore and Molecular Features to Survival Prediction in Stage III Colon Cancer. JNCI Cancer Spectrum, 4(3), pkaa023. 10.1093/jncics/pkaa023

Stefanescu, C., Van Gogh, M., Roblek, M., Heikenwalder, M., & Borsig, L. (2021). TGFβ Signaling in Myeloid Cells Promotes Lung and Liver Metastasis Through Different Mechanisms. Frontiers in Oncology, 11. https://www.frontiersin.org/articles/10.3389/fonc.2021.765151

Subramanian, A., Tamayo, P., Mootha, V. K., Mukherjee, S., Ebert, B. L., Gillette, M. A., Paulovich, A., Pomeroy, S. L., Golub, T. R., Lander, E. S., & Mesirov, J. P. (2005). Gene set enrichment analysis: A knowledge-based approach for interpreting genome-wide expression profiles. Proceedings of the National Academy of Sciences, 102(43), 15545– 15550. 10.1073/pnas.0506580102

Tecalco-Cruz, A. C., Sosa-Garrocho, M., Vázquez-Victorio, G., Ortiz-García, L., Domínguez-Hüttinger, E., & Macías-Silva, M. (2012). Transforming Growth Factor-β/SMAD Target Gene SKIL Is Negatively Regulated by the Transcriptional Cofactor Complex SNON-SMAD4. The Journal of Biological Chemistry, 287(32), 26764–26776. 10.1074/jbc.M112.386599

Therneau, T. M., & Grambsch, P. M. (2000). Modeling Survival Data: Extending the Cox Model. Springer. 10.1007/978-1-4757-3294-8

Thorsson, V., Gibbs, D. L., Brown, S. D., Wolf, D., Bortone, D. S., Ou Yang, T.-H., Porta-Pardo, E., Gao, G. F., Plaisier, C. L., Eddy, J. A., Ziv, E., Culhane, A. C., Paull, E. O., Sivakumar, I. K. A., Gentles, A. J., Malhotra, R., Farshidfar, F., Colaprico, A., Parker, J. S., … Shmulevich, I. (2018). The Immune Landscape of Cancer. Immunity, 48(4), 812–830.e14. 10.1016/j.immuni.2018.03.023

Tschernia, N. P., & Gulley, J. L. (2022). Tumor in the Crossfire: Inhibiting TGF-β to Enhance Cancer Immunotherapy. BioDrugs, 36(2), 153–180. 10.1007/s40259-022-00521-1

Tukey, J. W. & others. (1977). Exploratory data analysis (Vol. 2). Reading, MA.

Turan, T., Kongpachith, S., Halliwill, K., Roelands, J., Hendrickx, W., Marincola, F. M., Hudson, T. J., Jacob, H. J., Bedognetti, D., Samayoa, J., & Ceccarelli, M. (2021). A balance score between immune stimulatory and suppressive microenvironments identifies mediators of tumour immunity and predicts pan-cancer survival. British Journal of Cancer, 124(4), Article 4. 10.1038/s41416-020-01145-4

Vasey, B., Clifton, D. A., Collins, G. S., Denniston, A. K., Faes, L., Geerts, B. F., Liu, X., Morgan, L., Watkinson, P., McCulloch, P., & The DECIDE-AI Steering Group. (2021). DECIDE-AI: New reporting guidelines to bridge the development-to-implementation gap in clinical artificial intelligence. Nature Medicine, 27(2), Article 2. 10.1038/s41591-021-01229-5

Vincent, B. G., Szustakowski, J. D., Doshi, P., Mason, M., Guinney, J., & Carbone, D. P. (2021). Pursuing Better Biomarkers for Immunotherapy Response in Cancer Through a Crowdsourced Data Challenge. JCO Precision Oncology, 5, 51–54. 10.1200/PO.20.00371

Waskom, M. (2021). seaborn: Statistical data visualization. Journal of Open Source Software, 6(60), 3021. 10.21105/joss.03021

Wickham, H. (2016). ggplot2: Elegant Graphics for Data Analysis. Springer-Verlag New York. https://ggplot2.tidyverse.org

Wilks, C., Zheng, S. C., Chen, F. Y., Charles, R., Solomon, B., Ling, J. P., Imada, E. L., Zhang, D., Joseph, L., Leek, J. T., Jaffe, A. E., Nellore, A., Collado-Torres, L., Hansen, K. D., & Langmead, B. (2021). recount3: Summaries and queries for large-scale RNA-seq expression and splicing. Genome Biology, 22(1), 323. 10.1186/s13059-021-02533-6

Yadav, R. K., Chae, S.-W., Kim, H.-R., & Chae, H. J. (2014). Endoplasmic Reticulum Stress and Cancer. Journal of Cancer Prevention, 19(2), 75–88. 10.15430/JCP.2014.19.2.75

Yang, M., Lin, C., Wang, Y., Chen, K., Zhang, H., & Li, W. (2022). Identification of a cytokine-dominated immunosuppressive class in squamous cell lung carcinoma with implications for immunotherapy resistance. Genome Medicine, 14(1), 72. 10.1186/s13073-022-01079-x

Zanetti, M., Xian, S., Dosset, M., & Carter, H. (2022). The Unfolded Protein Response at the Tumor-Immune Interface. Frontiers in Immunology, 13, 823157. 10.3389/fimmu.2022.823157

Zhang, H., Klareskog, L., Matussek, A., Pfister, S. M., & Benson, M. (2019). Translating genomic medicine to the clinic: Challenges and opportunities. Genome Medicine, 11(1), 9. 10.1186/s13073-019-0622-1

Zhao, Y., Li, M.-C., Konaté, M. M., Chen, L., Das, B., Karlovich, C., Williams, P. M., Evrard, Y. A., Doroshow, J. H., & McShane, L. M. (2021). TPM, FPKM, or Normalized Counts? A Comparative Study of Quantification Measures for the Analysis of RNA-seq Data from the NCI Patient-Derived Models Repository. Journal of Translational Medicine, 19(1), 269. 10.1186/s12967-021-02936-w

